# URMAP, an ultra-fast read mapper

**DOI:** 10.1101/2020.01.12.903351

**Authors:** Robert C. Edgar

**Affiliations:** Independent Investigator, Corte Madera, California, USA

## Abstract

Mapping of reads to reference sequences is an essential step in a wide range of biological studies. The large size of datasets generated with next-generation sequencing technologies motivates the development of fast mapping software. Here, I describe URMAP, a new read mapping algorithm. URMAP is an order of magnitude faster than BWA and Bowtie2 with comparable accuracy on a benchmark test using simulated paired 150nt reads of a well-studied human genome. Software is freely available at https://drive5.com/urmap.

## Introduction

### Background

Next-generation sequencing has enabled dramatic advances in fields ranging from human functional genomics (Morozova and Marra, 2008) to microbial metagenomics (Gilbert and Dupont, 2011). Data analysis in next-generation studies often requires mapping of reads to a reference database such as a human genome, human exome, or a collection of full-length microbial genomes. Mapping is a special case of sequence database search where the query sequence is short, database sequences are long, and sequence similarity is high. For a given query sequence (read), the primary goal of mapping is to report the best match if possible, otherwise to report that the best two or more alignments are sufficiently similar to each other that the best match is ambiguous.

### Prior work

Many mapping algorithms have been proposed. Representative examples include BWA (Li and Durbin, 2009), Bowtie (Langmead *et al.*, 2009), Bowtie2 (Langmead and Salzberg, 2012), SOAP (Li *et al.*, 2008), SOAP2 (Li *et al.*, 2009b), Minimap2 (Li, 2018), FSVA (Liu *et al.*, 2016), SSAHA (Ning *et al.*, 2001) and SNAP (Zaharia *et al.*, 2011). Mappers utilizing the Burrows-Wheeler Transform (BWT) (Burrows and Wheeler, 1994) are the current *de facto* standard, with BWA and Bowtie2 in particular having more than 39,000 citations combined at the time of writing (Google Scholar accessed 31st Dec 2019). When first utilized in read mapping, BWT had the important advantage that it creates a compact index with size comparable to the reference database. For the human genome, this is ∼3Gb, which is small enough to be stored in RAM with the commodity computers of that time. Currently, computers with 32Gb or more RAM are readily available, which has raised the question of whether additional memory could enable better mapping performance. In particular, the authors of SNAP claim (Zaharia *et al.*, 2011) that its use of a ∼27Gb hash table index for the human genome gives both faster speed and higher accuracy than BWA. FSVA also uses a hash table, reportedly (Liu *et al.*, 2016) achieving faster speed than BWT though with somewhat lower accuracy.

### URMAP algorithm

URMAP uses a hash table index on *k*-mers, i.e. fixed-length words of length *k*, where *k*=24 is recommended for the human genome. The index is designed to keep information relating to a given hashed word (*slot*) close together in RAM to minimize memory cache misses. Slots found exactly once in the reference (*pins*) are flagged. For a given query, URMAP first searches for a pair of non-overlapping pins which are close together in the reference (a *brace*, see Fig. 1). If a brace is found, an alignment is attempted and the search terminates immediately if successful. Otherwise, a seed-and-extend strategy (Altschul *et al.*, 1990) is followed which prioritizes low-abundance slots.

**Figure 1.**
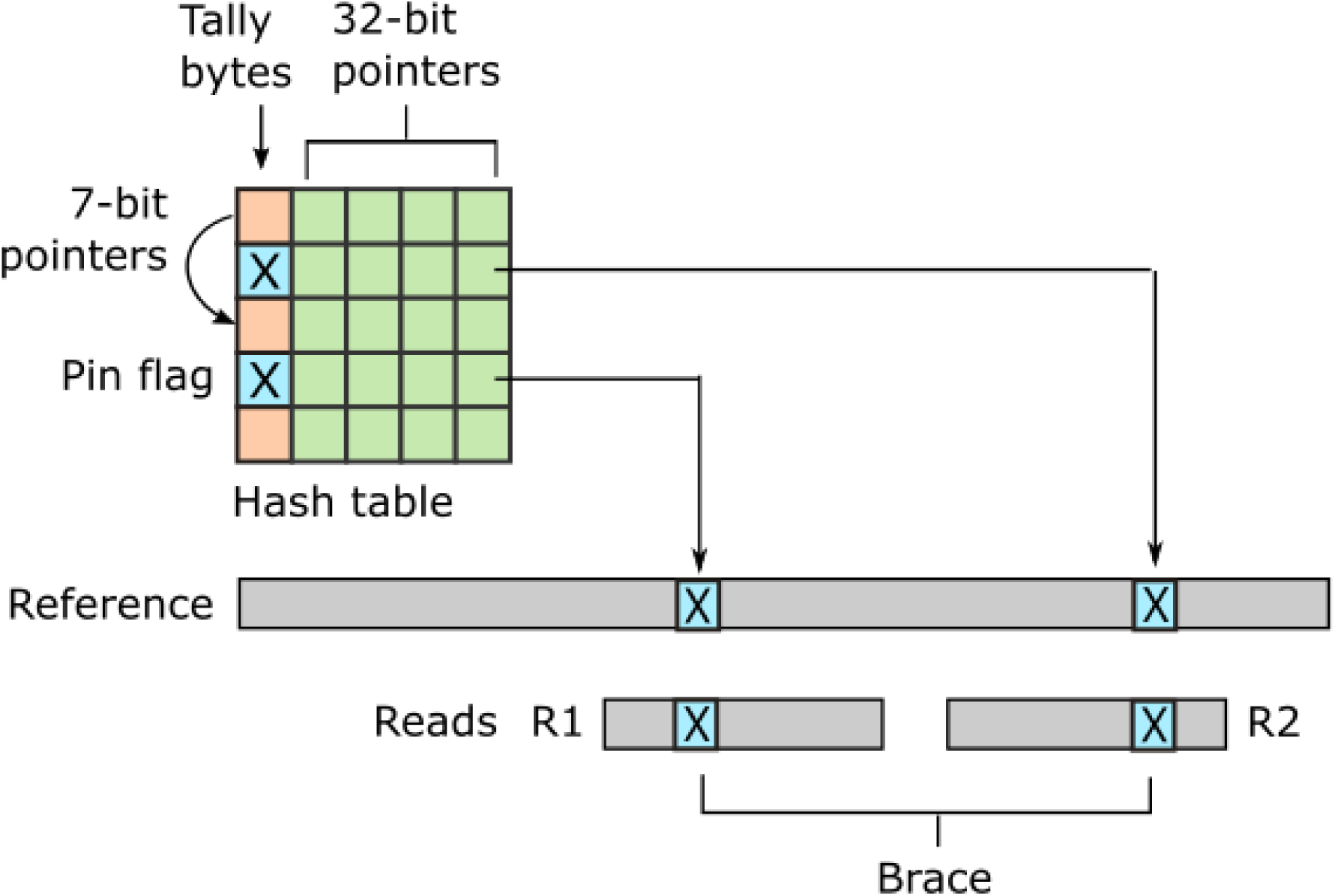
Schematic of the URMAP algorithm. Words in the plus strand of the reference sequence are indexed using a hash table with 5 bytes per row comprising a tally byte and a 32-bit pointer. Pins, i.e. words with a hash value that is unique across both strands of the reference, are indicated by a reserved tally value (pin flag). URMAP searches for a brace, i.e. a pair of pins close in the reference, one in the forward read (R1) and one in the reverse read (R2). If a brace is found, it is almost certain to be the correct location. Words found more than once in the reference are indexed using a linked list with forward pointers which are stored in tally bytes if they fit into 7 bits, otherwise in the 32-bit pointer field. The first bit of the tally is set if the row is in a list but not the head.

### Performance testing

Recent assessments of mapping accuracy, in particular those of SNAP and FSVA, have used the wgsim program in the SAMtools package (Li *et al.*, 2009a) to simulate reads of a human genome. Mutation rates (more correctly, variation rates) of 0.1% were used in both cases, with 0.09% single-nucleotide polymorphisms (SNPs) and 0.01% indels. The base call error rate was set to 0.4% for testing FSVA and to various different values for testing SNAP. Differences, i.e. base call errors, SNPs and indels, are introduced by wgsim with equal probability for each type at each position, giving a Poisson distribution for inter-difference spacing where closely-spaced SNPs and base call errors are rare. With a mutation rate of 0.1%, most reads of length 150nt simulated by wgsim have no mutation, and most reads with mutations have exactly one single-base variant. In real human genomes, variants tend to cluster, e.g. in non-coding regions (Altshuler *et al.*, 2010; Montgomery *et al.*, 2013). Thus, average accuracy over all reads on a wgsim test gives little insight into mapper performance on the more challenging, and often more biologically interesting, reads with multiple differences compared to the reference. Ilumina base call errors also tend to cluster, for example towards the end of a read (Minoche *et al.*, 2011), and in practice there are therefore many more reads with multiple errors than a Poisson distribution would predict.

### Urbench performance test

In this work, I introduce Urbench, a new benchmark test using experimentally determined variation from a well-characterized human genome. Simulated read sequences are combined with quality scores from a recent 2×150 Illumina run. At each base, a substitution error is introduced with the probability implied by its quality score, with the goal of generating a more realistic distribution of base call errors compared to earlier benchmarks. Mapping sensitivity and error rates are measured separately on reads which do, or do not, contain variants. Systematic errors are identified where most reads of a given locus are mapped to the same incorrect locus. Systematic errors are presumably more likely to cause false-positive inferences in later analysis than errors spread over many incorrect positions.

## Methods

### URMAP index

#### Hash table

The positions of words of length *k* in the plus strand of the reference are stored in a hash table. A word *W* is converted to an integer *w*(*W*) ∈ [0, 4^*k*^) in the usual way by considering letters to be base-4 digits `A`=0, `C`=1, `G`=2 and `T`=3. The `murmur64` hash function (https://en.wikipedia.org/wiki/MurmurHash) is used to convert *w* to an integer (*slot*) *s* ∈ [0, *H*) where *H* is the table size by *s* = `murmur64`(*w*(*W*)) `mod` *H*. For brevity, a word with a given slot value will be referred to simply as a slot. To reduce collisions, the table size *H* should be a prime number substantially larger than the reference; for the human genome *H*=5×10^9^ + 29 is recommended. The table design is intended to minimize size and optimize adjacency of data relating to a given slot with the goal of avoiding memory cache misses.

#### Hash table row

Each hash table entry (*row*) is five bytes: one byte (the *tally*) containing a one-bit flag and sometimes a 7-bit pointer to another row, and a four-byte value which may be a 32-bit reference coordinate or a pointer to another row. Using 32-bit coordinates limits the total reference sequence size to 2^32^ = 4 GB; references up to 1024 GB could be accommodated by using five pointer bytes.

#### Pins

A *pin* is a slot found exactly once in the reference, considering both plus and minus strands. This is an important special case because if a pin is found in a read it probably maps correctly to the same pin in the reference, though it may also be a false positive due to sequencing error or a genome variant. A pin is indicated by a reserved tally value. The term “pin” was chosen by analogy with a metal (or virtual) pin used to mark a location on a paper (or online) geographical map.

#### Singletons

A *singleton* is a slot that is found exactly once in the plus strand of the reference and one or more times in the minus strand. A singleton is also indicated by a reserved tally value. Note that while the minus strand is not indexed, there is nevertheless an important distinction between pins and singletons because the reverse-complement of a singleton occurs at least once in the plus strand of the reference, while the reverse-complement of a pin does not occur. Thus, while a pin found in a read implies only one candidate alignment to the plus strand of the reference considering both strands of the query, a singleton implies at least two candidates.

#### Linked lists

To store positions of a slot occurring more than once in the plus strand of the reference, a linked list is stored in nearby empty rows. Where possible, 7 bits of the tally are a pointer to the next row in the list, represented as the number of rows to skip. Overflows where this number does not fit into 7 bits are handled by storing a pointer to the next row in the 32-bit value instead of a reference coordinate. Overflows and list ends are indicated by reserved tally values.

#### Over-abundant slots

Slots exceeding an abundance threshold are excluded from the index. This is a speed optimization to avoid constructing a large number of candidate alignments. The loss in sensitivity is small because abundant slots are usually found in repetitive sequence which maps ambiguously unless there is distinctive sequence elsewhere in the read, and unique reference sequence of length ≥*k* necessarily contains a pin. By default, the abundance threshold *t* is set to 32. A slot that occurs more than *t* times on either reference strand is excluded. The minus strand is also considered in order to exclude cases where a repeat occurs with high abundance on the minus strand but low abundance (in particular, only once) on the plus strand. If a slot in such a repeat were indexed, this would tend to lead to an over-estimate of the probability that one of its plus strand alignments is correct because high-scoring secondary alignments to the minus strand would not be discovered.

#### Absent slots

A slot that does not occur in the reference, or is not indexed because it is over-abundant, is *absent*, as opposed to a slot which is present in the index. An absent slot may appear in a read, and the index must therefore indicate that the corresponding row does not contain a reference coordinate for that slot. This is accomplished by the first bit of the tally, which is set to one for present slots and zero for absent slots. The reserved tally values for pins and singletons have the first bit set to one to indicate that these slots are present, and the reserved values for 7-bit pointer overflows and the end of a linked list start with a zero bit because these slots were found to be absent and the corresponding rows were therefore available for use in a list. Linked list pointers in tally bytes are limited to 7 bits to ensure that the first bit is zero.

#### Collisions

A hash table collision occurs when two different reference words have the same slot value. A collision between the query and index occurs when a word in the read is different from a word in the index and has the same slot value. Collisions are not represented in the index or explicitly checked during search. This strategy saves index space without compromising search time because in the rare cases where a collided slot is aligned, the alignment will be abandoned quickly due to excessive mismatches in the flanking reference sequence. When aligning, it is faster to check the flanking sequence first than to verify that the seed matches because in the typical (non-collision) case the seed always matches while flanking sequence often does not.

#### Word length

Increasing *k* increases the frequency of pins in the reference and also increases the number of words per query that are changed by a difference and hence the probability that a read does not contain a pin, or any indexed slot, due to read errors and variants. The choice of *k* is thus a compromise between speed and sensitivity. With the human genome, *k*=24 is recommended because of the ∼3G 24-mer slots, ∼0.9G (30%) are pins, and on average a reference segment of length 150nt contains 38 pins. A query sequence with <7 single-base differences is guaranteed to have at least one 24-mer match, noting that 7 differences eliminate all 24-mer matches only in the tiny fraction of possible distributions where they are maximally disruptive, and reads with ≥7 differences will often have at least one preserved 24-mer.

### URMAP search algorithm

#### Query word search order

With 24-mers, query words in a read of length 150 are processed at intervals (*strides*) of length 29 using modulo 127 to keep the position within the read (because there are 127 24-mers in a read of length 150). For example, the first three words processed are at positions 0, 29 and 58. At a given position, both strands are considered, so for example the words at the first position (zero) in both plus and minus strands are both processed before moving on to the plus and minus words at position 29. The stride value 29 is chosen to be relatively prime with 24, which ensures that the following loop will visit each query word exactly once:

~~~
for (int j = 0; j < 127; ++j) {
   QueryWordPosition = (29*j)%127;
   /* … */}.
~~~

The simple form of this loop without conditional branches may enable loop unrolling or vector parallelization by the compiler, and regardless is designed to be efficiently executed on modern processors. In general, given the word length *k*, the stride is identified as the smallest prime number with value ≥*k*+5. The use of a stride >*k* is motivated by the observation that neighboring words are not independent. If a query word is not a pin, fails to align, or is not indexed, this is likely to be because the word is in a repetitive region or variant, or contains a sequencing error. The immediately following words are likely to have the same problem, and the chances of finding good alignments early in the search are improved by skipping ahead.

#### First pass: brace search

In its first pass through the query words, URMAP seeks a pair of pins that are close together in the reference. As noted in the Introduction, such a pair is called a brace (Fig. 1). This term was chosen because the noun “brace” has two relevant meanings: two of a kind, and a device that connects, fastens or stabilizes. With paired reads, one pin is sought in the forward read (R1) and the other in the reverse read (R2). While a pin may be a false positive due to sequencing error or a variant, a brace is almost certain to be a true positive match. If a brace is found and is aligned successfully (is a *good brace*), the search terminates immediately. Previous read mapping algorithms do not terminate when the first high-scoring alignment is found, even if it has no mismatches, because equally high-scoring alignments may exist elsewhere in the reference. By contrast, when a good brace is found the likelihood that a different position is correct is vanishingly small. Noting that a typical read contains several pins, and most pins are true positives, the search for a good brace in a read pair proceeds as follows, with the goal of minimizing the number of hash table accesses and attempted alignments in typical cases. The first pins in both reads (the forward and reverse read, known as R1 and R2 respectively) are identified. If this pair is not a good brace, the next pin is identified in R1, giving a new potential brace, then the next pin in R2, and so on. This process continues until a good brace is found or all words in both reads have been processed. Almost all pin pairs which are not braces can be Identified as such because they are too far apart in the reference, which requires only the coordinates in their hash table rows. It is very rare for non-overlapping false positive pins to appear close in the reference, and therefore brace tests almost never fail in the more expensive alignment stage. Since most human 2×150 read pairs contain a correct brace, the brace search pass identifies the correct reference coordinate for most reads with remarkable efficiency.

#### Second pass: low-abundance slot search

If no brace is found, the hash table row for each query word has been accessed exactly once. Each row access almost certainly triggers a memory cache miss because of the large size of the hash table (∼25 GB for the human genome). To accelerate access to these rows in subsequent passes, the first pass copies them to a small (few kB) per-thread buffer (PTB). Other data which may be used repeatedly, such as query slot values and the reverse-complemented query sequence, is also stored in the PTB, which is designed to be compact and contiguous to maximize the chance that it will be available in a fast memory cache. The second pass attempts to align all non-pin slots with abundance ≤2. Most of the index data needed for this task is already present in the PTB, though some additional rows may be required for slots with abundance two. If a high-scoring alignment is found in the second pass, the search terminates.

#### Third pass: high-abundance slot search

In the rare case that no high scoring alignment is found in the first two passes, alignments are attempted for the remaining slots.

#### HSP construction

Following BLAST (Altschul *et al.*, 1990) and many subsequent algorithms, URMAP constructs ungapped alignments using a seed-and-extend strategy. The seed is an indexed slot found in the query, which implies an alignment of length *k*. The seed is extended into flanking sequence using gapless *x*-drop alignment which stops if the score falls more than *x* below the maximum so far observed (*x* is a heuristic parameter). If the score exceeds a threshold, the alignment is designated a high-scoring segment pair (HSP) and stored, otherwise the reference location is added to a list of failed extensions. The lists of HSPs and failed locations are consulted before extending to prevent redundant attempts to align the same reference location. If the HSP covers the entire query sequence, then the alignment is considered successful.

#### Gapped alignments

In the human genome, indel variants are rare (Altshuler *et al.*, 2010), and Illumina indel errors are very rare (Schirmer *et al.*, 2015), and therefore a large majority of correct alignments of human reads are expected to be gapless. Computing an ungapped alignment is much faster than a gapped alignment, and URMAP therefore constructs gapped alignments only if no HSP covers the query. Gapped alignments are constructed by extending the top few HSPs into semi-global alignments using a variant of the Viterbi algorithm (Viterbi, 2006) where the alignment is constrained to include the HSP and the terminal regions are banded, greatly reducing the number of dynamic programming matrix cells which must be computed. Here, semi-global means that the entire query sequence must be included but not the entire reference.

#### MAPQ calculation

MAPQ is an integer value representing the estimated probability *P*_*error*_ that the reference coordinate of the top-scoring alignment is wrong,

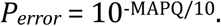

Let *T* be the score of the first alignment in order of decreasing score, and *S* ≤ *T* be the score of the second alignment. If only one alignment is found, *S* is set to *T*/2 as a prior estimate of the second-best score rather than zero because the URMAP search algorithm may terminate early if a high-scoring alignment is found. The first alignment is likely to be correct (*P*_*error*_ is small) if *T* ≫ *S*, and conversely *Perror* is at least ∼0.5 if *T* ≈*S* (because if exactly two alignments *X* and *Y* have equal scores, there is a 1/2 chance that *X* is wrong; 2/3 chance if there are three, and so on). Thus MAPQ should increase monotonically with *T - S*. Also, MAPQ should decrease monotonically with decreasing *T* because alignments with more differences are less likely to be correct. The best possible alignment score is the query sequence length |*Q*| because identities contribute 1 to the score, and the ratio *T*/|*Q*| therefore ranges from one to zero as the top alignment score ranges from best possible to worst possible. This ratio is a natural choice to down-weight *T* - *S*, and a simple formula with the desired properties is MAPQ = (*T-S*) *T*/|*Q*|. Empirically, using (*T*/|*Q*|)^2^ rather than *T*/|*Q*| was found to give a more accurate estimate, and URMAP therefore uses

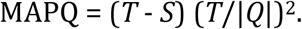

I am aware of no justification for this formula beyond its empirical success and the intuitive considerations above. However, the simplicity of this formula and its lack of tuned parameters (except perhaps the power of *T/*|*Q*|) suggest that this or a similar result may be derivable from more rigorous theoretical considerations.

### URMAPv algorithm

Some applications are more tolerant of mapping errors than variant calling, for example nucleosome position inference in cell-free DNA (Snyder *et al.*, 2016). With this in mind, I sought a set of parameters for URMAP giving faster execution time while maintaining useful accuracy. The most important speed optimization is reducing the maximum indexed slot abundance *t* from 32 to 3. Other optimizations include tweaks to heuristic parameters which trigger early termination of various search stages, such as the *x* in *x*-drop. Here this algorithm is called URMAPv; it is invoked by the *-veryfast* command-line option. In practice, the execution time of URMAPv is often dominated by file i/o, and thus represents a point of diminishing returns in speed optimization for mapping.

### Urbench benchmark test

I implemented a benchmark, Urbench, which models mapping of shotgun 2×150 Illumina reads, i.e. the current *de facto* standard, to the human reference genome. Variants and sequencing error were introduced based on experimental results rather than by simulating Poisson distributions as in previous benchmarks.

#### Reference sequence

Genome Reference Consortium Human genome build 38 (GRCh38) (Church *et al.*, 2011) was used as the reference sequence.

#### Genome variants

Experimentally determined variants with their coordinates relative to GRCh38 were taken from NA12878, a well-studied diploid human genome (Zhang *et al.*, 2019). Many of these variants are phased, i.e. assigned to a parental chromosome, over regions of tens to hundreds of kb. Unphased variants were randomly assigned to a parental chromosome.

#### Simulated read pairs

Two source genome sequences were used to simulate reads: the reference genome GRCh38 and the diploid variant genome NA12878. Using the reference models a situation where a true read sequence is identical to the reference (i.e., does not contain a variant), which is probably the case for most reads in practice. For each genome, 1M loci were selected. For each locus, ten read pairs were simulated at random positions such that R1 or R2 contained the locus (Fig. 2). This enables systematic errors to be identified, i.e. cases where the majority of reads for a given locus are assigned to the same incorrect reference segment. With NA12878, each locus was the position of a variant so that all simulated read pairs contains at least one variant. With GRCh38, loci were randomly selected positions.

**Figure 2.**
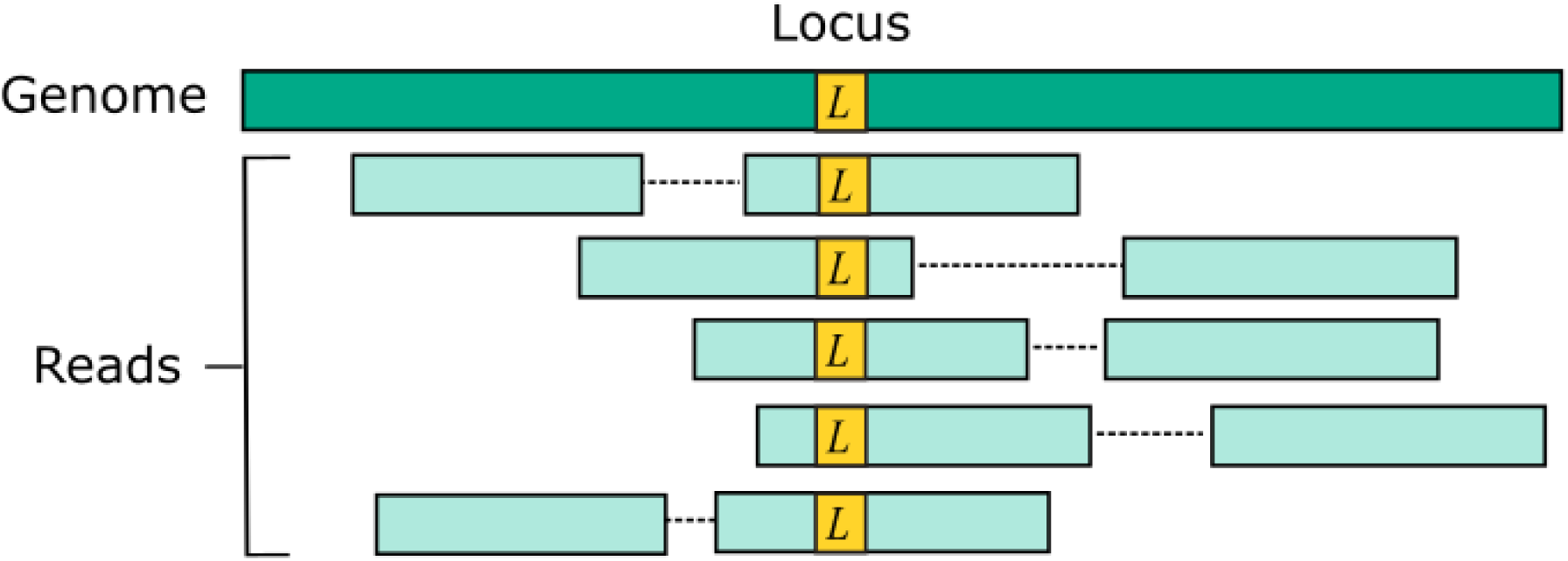
Design of the Urbench benchmark. For each locus *L* in a source genome (NA12878 or GRCh38), ten simulated reads pairs are generated (five shown in figure) such that either R1 or R2 contains the locus. This enables systematic errors to be identified where a majority of reads of a given locus are mapped to the same incorrect location. With NA12878 a locus is the position of an experimentally determined variant (SNP or indel) in one of the parental chromosomes, with GRCh38 a locus is a randomly-chosen position. Base call substitution errors are introduced with probabilities given by quality scores in sequencing run SRR9091899.

#### Sequencing error

Simulated read sequences were combined with quality scores from run SRR9091899 in the Sequence Read Archive (Leinonen *et al.*, 2011), which is a recent (submitted 2019) 2×150 Illumina shotgun dataset. At each base, a substitution error was introduced into the nucleotide sequence with the probability implied by its quality score.

#### Accuracy metrics

Per-read sensitivity *S*_*r*_ is defined as the fraction of reads which are mapped to the correct coordinate with high confidence as reported by the mapper. Following the BWA paper, high confidence was determined as MAPQ≥10, corresponding to *P*_*error*_≤0.1. Per-read error *E*_*r*_ is defined the fraction of reads with MAPQ≥10 which are mapped to an incorrect position. Per-locus sensitivity *S*_*l*_ is defined as the fraction of loci where at least three reads have MAPQ≥10 and the majority of these are mapped to the correct position. Per-locus error rate *E*_*l*_ is defined as the fraction of loci with at least three reads having MAPQ≥10 where the majority of these are mapped to the same incorrect position. *E*_*l*_ is interpreted as assessing systematic errors that are more likely to be harmful to downstream analysis than randomly-distributed errors. These metrics are measured separately for reads of the reference (*ref*) and of the variant genome (*var*) giving a total of eight accuracy metrics *S*^*ref*^_*r*_, *E*^*ref*^_*r*_, *S*^*ref*^_*l*_, *E*^*ref*^_*l*_, *S*^*var*^_*r*_, *E*^*var*^_*r*_, *S*^*var*^_*l*_ and *E*^*var*^_*l*_ which are expressed as percentages.

#### Pairwise method comparison

To enable a compact summary comparison of the eight accuracy metrics for a pair of methods *X* and *Y*, I defined the *mean improvement* of *X* over *Y* (MI_*XY*_) as the mean of *S*_*X*_ - *S*_*Y*_ - *E*_*X*_ + *E*_*Y*_ over the four combinations of genome (*ref* or *var*) and read or locus (*r, l*) and the *improved metric count* (IM) as the number out of the eight metrics where *X* has a better value than *Y* (higher if sensitivity, lower if error rate). If IM_*XY*_ is 8, then all the metrics for *X* are better than *Y*, and *X* is unambiguously better than *Y* by the Urbench test, denoted *X*>>*Y*. Conversely if IM_*XY*_ is zero then all metrics for *X* are worse, MI_*XY*_ is negative, and *X* is worse than *Y*, denoted *X*<<*Y*. If six out of eight *X* metrics are better; this is denoted by X>6*Y*, if five out of eight *X* metrics are worse, this is written *X*<5*Y*. The magnitude of the improvement is indicated by MI and written in parentheses, e.g. *X*<<(−1.2)*Y* or *X*>6(4.0)*Y*. The total improvement (TI_*X*_) of method *X* over the other tested methods is calculated as the total of MI over pairwise comparisons with other methods.

#### Speed

The time required to map a given set of reads depends on the computer architecture (e.g., the processor type, number of cores, and sizes and speeds of L1 and L2 memory caches) and overhead due to file input/output (i/o). Reads are typically provided in compressed FASTQ format (fastq.gz extension), which requires potentially expensive decompression, and output is typically written to large SAM (uncompressed) or BAM (compressed) files. The overhead of file i/o (including decompression and/or compression, if applicable) can be substantial for the faster mappers and varies widely with the computing environment. With this in mind, I chose to measure speed using a method designed to isolate mapping by reducing i/o overhead as much as possible, as follows. 10M reads were selected at random from SRR9091899, giving a total of ∼2.4 GB compressed (∼6GB uncompressed) FASTQ data (∼1.2 GB / 3Gb each for R1 and R2). These files are small enough to be cached in memory by the operating system, which minimizes i/o time on a computer with sufficiently large RAM. Using a PC with a 16-core Intel i7-7820X CPU and 64 GB RAM (more than twice the size of the largest genome index), I ran each mapper three times in succession using from 12 to 20 threads, first with uncompressed then compressed reads, and selected the shortest time after subtracting the time required to load the genome index. Speed is expressed as a multiple of the shortest time for BWA, i.e. the speed of BWA is 1.0 by definition. With this method, the measured relative speed of two mappers can be interpreted as a limit on the ratio in practice which would be approached by the fastest possible i/o.

#### Accuracy of MAPQ

The accuracy of MAPQ was assessed as follows. For each integer value (*q*) of MAPQ reported by a mapper, the total number of reads *n*^*q*^ and number of incorrectly mapped reads *n*^*q*^_*error*_ were calculated, giving the measured error frequency *f*^*q*^_*error*_ = *n*^*q*^_*error*_ / *n*^*q*^ and the measured mapping quality at the reported quality *q* is then

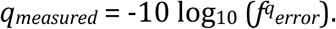

If the MAPQ values are accurate, then *q*_*measured*_ should be ∼*q* for all reported values of *q*, which was assessed by constructing a scatterplot of *q*_*measured*_ against *q*.

#### Tested methods

The following methods were tested: BWA v0.7.17-r1188, Bowtie2 v2.3.4.1, SNAP v1.0beta.24, FSVA GitHub commit 8cec132 (dated Jul 29, 2016), Minimap2 v2.17-r94 and URMAP v1.0.1300. This is a somewhat arbitrary selection from the many published methods designed include the most popular software (BWA and Bowtie2) together with potentially competitive recently-published methods. The beta version of SNAP was used because the release binaries failed on some tests.

## Results

### Speed and accuracy on Urbench

Metrics for speed and accuracy are shown in Table 1, with pair-wise summary comparisons in Table 2. As with most benchmarks in biological sequence analysis, results should be interpreted with caution because of the limitations of simulated data and the many somewhat arbitrary decisions that must be made in designing the benchmark and its performance metrics; other defensible designs would no doubt give somewhat different method rankings and numerical values for sensitivity and error rates. With these caveats in mind, some general trends can be observed from Tables 1 and 2. URMAP and URMAPv are the fastest methods, with URMAP ∼9× faster than BWA and Bowtie and URMAPv ∼20× faster, noting that in practice the speed improvement may be less due to file i/o overhead. All methods were faster with uncompressed FASTQ, showing that the added time for decompression exceeds the time saved by reading smaller files. Three methods, BWA, URMAP and Bowtie2, stand out as more accurate than the others (SNAP, Minimap2, URMAPv and FSVA) because all methods from the first group have at least 6 better metrics (shown as >6, >7 or >> in Table 2) with a positive mean improvement compared to all methods in the second group. In particular, BWA is unambiguously more accurate than SNAP as all eight metrics are better with a mean improvement of 3.2, contradicting the claim of the SNAP authors that it is more accurate than BWA. The three better methods have similar accuracy, with no method having more than 5 out of 8 better (hence 3 or 4 worse) accuracy metrics than another, and all pairwise comparisons exhibit only small mean improvements ranging from BWA >5(0.5) URMAP to BWA >5(2.0) Bowtie2. Thus, the accuracy differences between BWA, URMAP and Bowtie2 are small and ambiguous, and I believe these differences are unlikely to be consequential in practice for most applications.

**Table 1.** Performance on Urbench. Speed is measured relative to BWA with file i/o overhead minimized. Accuracy metrics are *S* (sensitivity) and *E* (error rate) with MAPQ≥10, expressed as percentages. Superscript is *var* for the variant genome (NA12878) or *ref* for the reference (GRCh38), subscript is *r* for per-read or *l* for per-locus.

| *Method* | *Speed* | $S^{var_r}$ | $E^{var_r}$ | $S^{var_l}$ | $E^{var_l}$ | $S^{ref_r}$ | $E^{ref_r}$ | $S^{ref_l}$ | $E^{ref_l}$ |
| --- | --- | --- | --- | --- | --- | --- | --- | --- | --- |
| Bowtie2 | 1.1 | 92.4 | 0.09 | 94.5 | 0.06 | 94.9 | 0.02 | 95.7 | 0.00 |
| BWA | 1.0 | 96.3 | 1.21 | 97.4 | 1.20 | 96.7 | 0.01 | 97.3 | 0.00 |
| FSVA | 4.6 | 88.9 | 3.79 | 94.0 | 2.20 | 93.8 | 1.23 | 96.1 | 0.11 |
| Minimap2 | 1.4 | 91.0 | 0.88 | 93.8 | 1.30 | 93.7 | 0.02 | 95.5 | 0.00 |
| SNAP | 6.4 | 92.3 | 2.04 | 94.1 | 2.00 | 94.8 | 0.15 | 95.6 | 0.05 |
| URMAP | 8.9 | 95.2 | 0.78 | 96.7 | 0.65 | 96.1 | 0.16 | 96.8 | 0.00 |
| URMAPv | 21 | 90.0 | 0.97 | 92.7 | 1.10 | 93.7 | 0.27 | 95.2 | 0.01 |

**Table 2.**
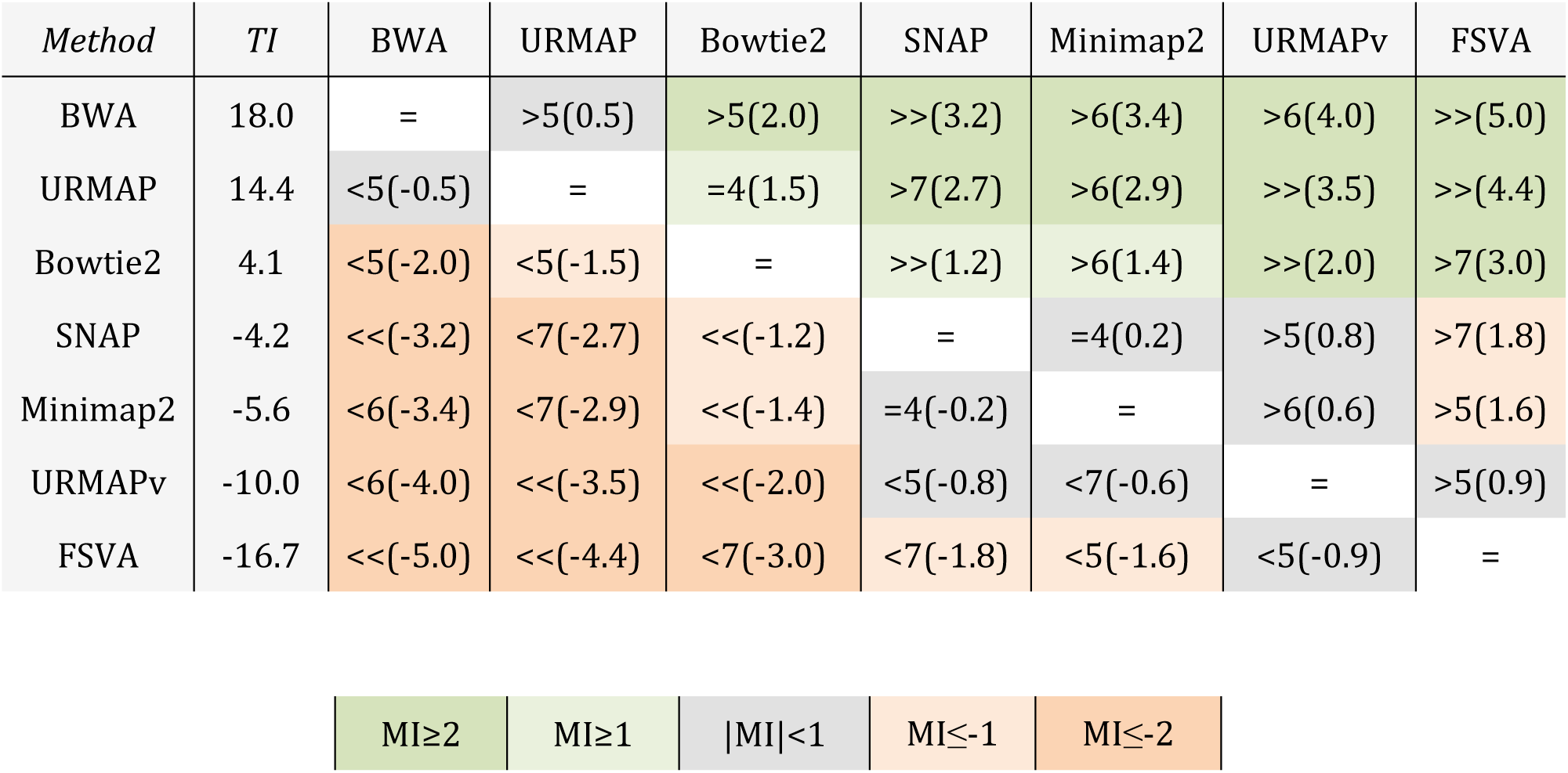
Pair-wise method comparisons on Urbench. Methods are sorted by decreasing total improvement (TI). Table cells are colored according to mean improvement (MI) as shown in the key. A pairwise comparison of the method in row *X* vs. the method in column *Y* is given using the notation described in Methods; e.g. BWA >5(2.0) Bowtie2 means that BWA has five of eight metrics that are better than Bowtie2 with a mean improvement of 2.0. The symbols >> and << indicate that all metrics are better or worse, respectively, e.g. BWA >>(3.2) SNAP means that BWA is better than SNAP by all metrics with a mean improvement of 3.2.

### MAPQ accuracy

Fig. 3 is a scatterplot of reported vs. measured MAPQ for the tested methods. Bowtie2 and URMAP are close to the diagonal, showing reasonably good estimates of MAPQ though Bowtie2 tends to underestimate and URMAP tends to overestimate. The other tested methods have much stronger tendencies to overestimate. For example, with MAPQ=50 (estimated *P*_*error*_ = 10^−5^), the measured MAPQ for BWA is 15.7 (*P*_*error*_ = 0.027) and the measured MAPQ for SNAP is 5.6 (*P*_*error*_ = 0.28).

**Figure 3.**
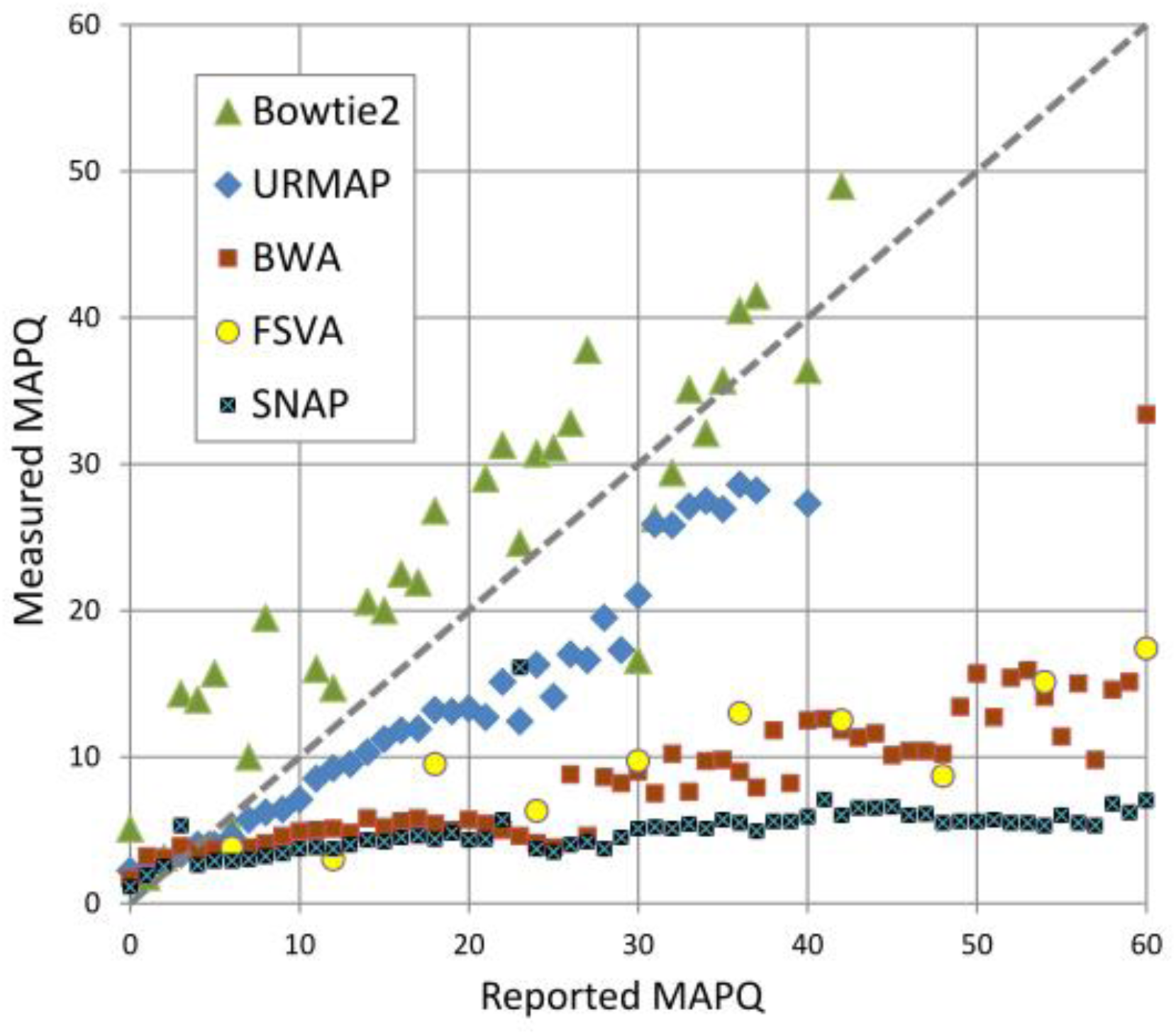
Scatterplot of reported MAPQ vs. measured MAPQ. For each integer value of MAPQ, the measured MAPQ is determined by the frequency of incorrectly mapped reads in the Urbench benchmark.

## Discussion

The results presented here show that URMAP is an order of magnitude faster than the popular BWT mappers BWA and Bowtie2 while achieving comparable accuracy. Other tested mappers using a hash table index, including SNAP, Minimap2 and FSVA, are less accurate. The quality estimates (MAPQ values) reported by URMAP and Bowtie2 are quite good, while BWA substantially overestimates quality, i.e. underestimates error probability. All mappers are faster with uncompressed FASTQ, which suggests that gzip is a poor choice of compression algorithm. Faster compression with similar size efficiency could be achieved by a FASTQ-specific algorithm, e.g. by using 2-bit values for the four nucleotide letters.

